# Morphology-Guided Deep Learning for Nanoparticle Agglomeration Diagnostic Assays

**DOI:** 10.1101/2025.09.13.676051

**Authors:** Kartik Jhawar, Xiao-Liu Chu, Joseph B. DeGrandchamp, Yueming Yin, Lalitha Anjali Peddibhotla, Shabina Banu, Farah Najwa Nabila Mohd Hatta, Yaoli Wang, Peter Török, Lipo Wang

**Affiliations:** Institute for Digital Molecular Analytics and Science, Nanyang Technological University, Singapore 636921; School of Electrical and Electronic Engineering, Nanyang Technological University, Singapore 639798; College of Computing and Data Science, Nanyang Technological University, Singapore 639798; Republic Polytechnic, Singapore 738964; Universiti Putra Malaysia, Selangor Darul Ehsan 43400, Malaysia; College of Electronic and Information Engineering, Taiyuan University of Technology, Taiyuan, China; Singapore Centre for Environmental Life Sciences Engineering, Nanyang Technological University, Singapore 637551; Lee Kong Chian School of Medicine, Nanyang Technological University, Singapore 636921; School of Physical and Mathematical Sciences, Nanyang Technological University, Singapore 637371

**Author notes:** These authors contributed equally to this work (Kartik Jhawar, Xiao-Liu Chu, Joseph B. DeGrandchamp).

## Abstract

Affordable, accurate, and rapid point-of-care diagnostic tests remain elusive due to inherent trade-offs between performance and cost. Conventional nucleic acid tests offer high sensitivity but require complex, expensive steps such as amplification and purification, whereas lateral flow assays are simple and low-cost but lack the necessary sensitivity for many applications. To bridge this gap, we present a miniaturized and simplified chip-based platform that combines three components into a single diagnostic pipeline: we use (i) spectrally distinct silver and gold nanoparticles that form analyte-dependent clusters with unique spectral fingerprints, (ii) a one-pot, enzyme- and purification-free assay on a chip integrated with a high-throughput automated low-cost microscope, and (iii) a morphology-guided convolutional Graph Neural Network that embeds morphology information into convolutional kernels and performs graph-based relational learning across particle-level features. This integration captures spectral, spatial, and morphological quantification at the particle level, rather than relying on bulk spectral shifts, thereby overcoming the limitations of contemporary nanoparticle assays and image-level deep learning approaches. Processing up to 5000 particles per image using only <5 GB GPU memory, Mc-GNN achieves femtomolar sensitivity with 98.2% recall for synthetic DNA and 94.8% for SARS-CoV-2 RNA from whole virus, despite variations in nanoparticle selection and sample complexity. By embedding morphological information into the biosensing pipeline, our diagnostic platform is computationally efficient, smartphone-compatible and is readily extensible to new analytes and multiplexing, offering a scalable solution for a fieldable diagnostic tool.

## 1 Introduction

The COVID-19 pandemic exposed the limits of infectious disease diagnostics. Nucleic acid amplification tests such as reverse transcription polymerase chain reaction (RT-PCR) remain highly sensitive but require costly equipment, long workflows, and centralized laboratories^1,2^. Antigen and serological assays are more accessible but often lose sensitivity and specificity at the early infection stage, when viral loads are low^2^. There is therefore a continuing need for rapid, affordable, and reliable point-of-care (POC) diagnostics that function with minimal infrastructure.

Lateral flow assays (LFAs) partially meet these criteria but miss low-abundance targets, which therefore yield false negatives^3,4^. Colorimetric nanoparticle (NP) assays promise higher sensitivity by exploiting analyte-driven aggregation and the resulting shifts in localized surface plasmon resonances (LSPRs). Foundational work has shown that DNA-induced gold NP aggregation can indicate particle cross-linking via distinct optical changes^5–7^. However, as is presented in Fig.1a, a red-shift in the plasmonic resonance is highly dependent on the interparticle distance. For a dimer comprising two gold NPs, a 40 nm separation produces only a 5 nm shift, which is challenging to detect without a high-end spectrometer. Only for interparticle spacings below 10 nm is the LSPR shift ≥30 nm, hence limiting this methodology to targets that bring NPs to close proximity and use high-end detectors.

To solve this challenge, several strategies have been explored. Spectrally distinct hetero-nanoparticle systems produce multi-colour signals upon target binding, avoiding the need for a large LSPR signal shift^7–9^. Other approaches combine NP assays with image-based readouts and machine learning on consumer optics^10–12^. However, these implementations typically depend on coarse bulk-colour features, manual counting, or simple classifiers, lacking particle-feature level insights essential for high sensitivity^13–18^. More recent deep learning (DL) efforts improve sensitivity but mainly operate at the whole-image level, lacking the deeper analyses at the particle-level, whilst requiring heavy computation^10,11,19–25^. A limited number of studies, including that of Shiaelis *et al*., have explored particle-resolved DL approaches, but in this case relying on fluorescently labelled viruses in total-internal-reflection fluorescence microscopy^21^. Here we present a simple, one-pot assay and analysis pipeline that integrates three components: (i) analyte-induced aggregation of spectrally distinct NPs, (ii) image-based single-NP detection on an automated dark-field microscope with low magnification, and (iii) a morphology-guided deep learning model (Mc-GNN) that extracts particle-level spectral, spatial, and morphological cues without fluorescent labels or region-of-interest (ROI) alignment. And in contrast to earlier hetero-particle implementations^7–9^, our method simplifies the assay into a one-pot, purification-free protocol. Notably, unlike standard nucleic acid tests such as RT-PCR^26,27^, this one-pot design is feasible because the detection chemistry is entirely protein-free, allowing for incorporation of lysing reagents directly for the release nucleic acid targets without inhibiting performance^28–30^. By incorporating our Mc-GNN architecture, we are able to robustly discriminate analyte concentrations down to the femtomolar regime, taking into account unbound particles and small clusters without the need for fluorescence labelling or ROI alignment^19,21^. We directly compare our DL method with other contemporary approaches and find that it outperforms them in terms of quantifying concentrations of DNA and SARS-CoV-2 viral RNA samples, demonstrating a high accuracy, computational efficiency, and assay robustness despite day-to-day sample variations. Our ablation study further confirms that the combination of incorporating the morphological information about nanoparticles and clusters with relational graph modelling is essential in achieving the high classification accuracy observed. By merging affordable optical imaging, NP spectral differentiation, and an intuitive DL architecture, our Mc-GNN-powered assay addresses longstanding limitations in POC diagnostics, providing a miniaturised, affordable, sensitive, and scalable solution that will unite laboratory performance with real-world diagnostic needs.

## 2 Results and Discussion

### 2.1 Numerical simulations

In nanoparticle aggregation assays, the main challenge is to distinguish single unbound particles from analyte-induced clusters. At low target concentrations, most particles remain unbound, in dimers or in small clusters, which are difficult to resolve by size or scattering intensity alone. Spectral cues provide a crucial additional channel, and we therefore employ gold-shell silica-core NPs (Au@SiNPs, 80 nm diameter silica + 20 nm thick shell) as red scatterers, 25 nm radius spherical silver NPs (AgNPs) as blue scatterers, and, in the final iteration, 25 nm radius spherical gold NPs (AuNPs) as green scatterers (see Supplementary Information (SI) (Figure S2)). We use the spectral profile of a single point spread function (PSF) to identify NP clustering types via co-localization. At low target concentrations, where high sensitivity is crucial, both specific and non-specific binding can form dimers. Bound NPs are typically separated by approximately 40 nm due to the size of the target molecule, whereas non-specific binding often results in smaller interparticle distances as nanoparticles simply stick together. To quantify the spectral shifts caused by dimer and trimer formation for different interparticle separations, we performed numerical simulations using COMSOL Multiphysics. Figure 1b and c demonstrate that the scattering contribution is predominantly from the larger Au@SiNPs. Although the magnitude of the LSPR spectral redshift is governed by the interparticle distance and shows less dependence on the second nanoparticle type (AgNPs or AuNPs), the presence of AgNPs introduces a distinctive spectral peak at approximately 450 nm. This peak decreases with reduced interparticle spacing when dimers form. Similar trends are demonstrated for trimers in Fig. 1d. As such, the ratio of blue, green and red colour intensities can provide measurable parameters to discriminate between specific and non-specific binding events through spectral analysis of the particle ensemble in an nanoparticle assay.

**Figure 1.**
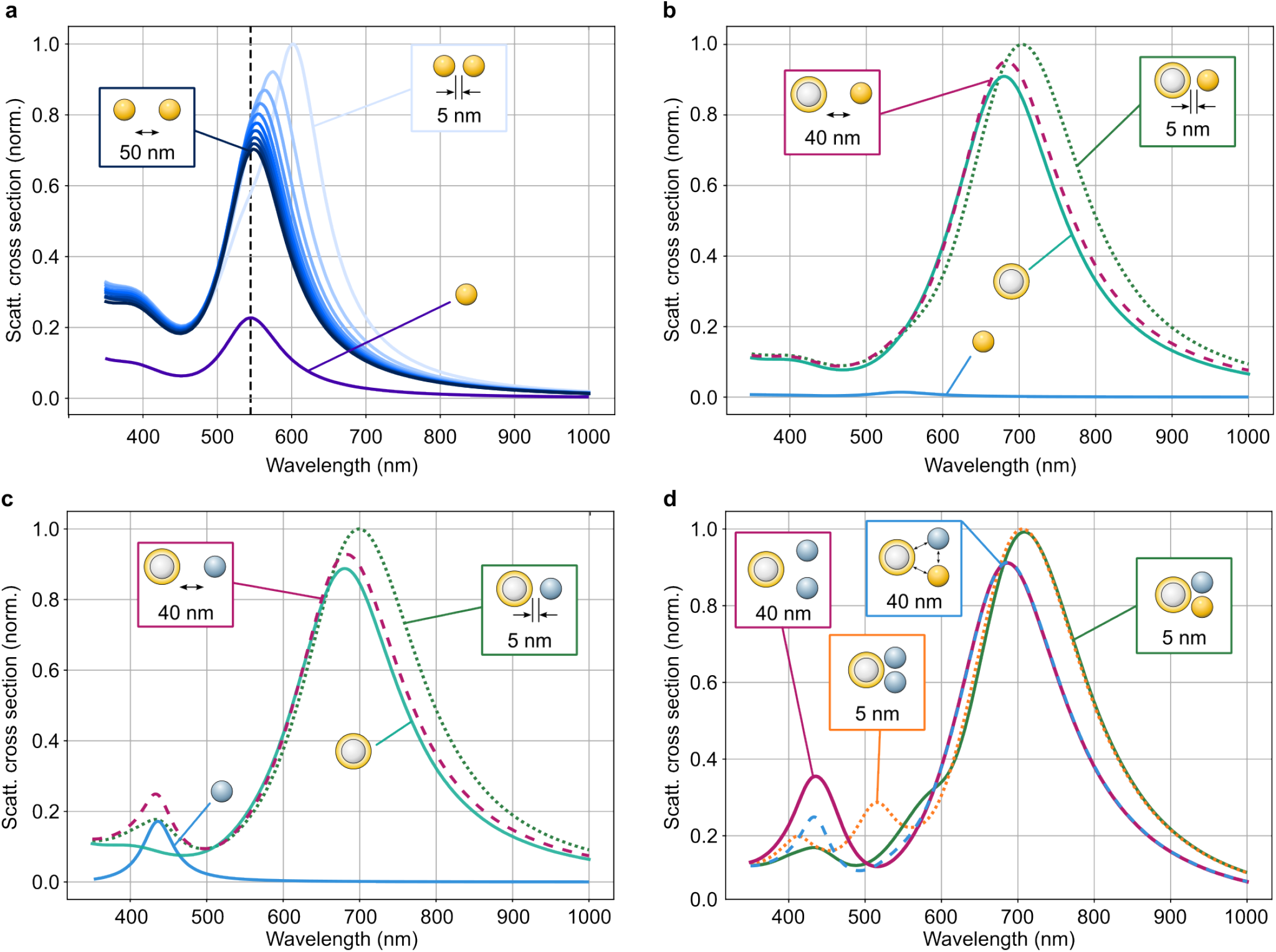
Numerical simulations of scattering cross sections of various nanoparticle assemblies. (a) The scattering cross section redshifts when the interparticle distance between two Au NPs decreases from 50 to 5 nm. The dashed line provides a guide for the eye for a single 25 nm radius Au NP. The scattering cross-section shows a similar trend for both a (b) Au@SiNP-AuNP dimer and a (c) Au@SiNP-AgNP dimer assembly. (d) Larger clusters, including Au@SiNP-AuNP-AgNP or Au@SiNP-2AgNP trimers, exhibit both distance-dependent redshifts and distance-dependent intensity variations of the AgNP spectral peak. The simulation is setup with unpolarized excitation propagating into the page (out-of-plane direction).

### 2.2 Overview of the experimental method

The experimental setup is shown in Figure 2, where we employed a home-built dark-field microscope to capture spectral signatures of nanoparticles. Broad-spectrum LEDs (Cree C503D-WAN, 40k mcd intensity) provided low-cost illumination, yielding measurable colour differences between clusters and single particles. Images that include this spectral information of different NP sub-populations was collected using a colour (RGB) camera (Teledyne FLIR Grasshopper 3, model GS3-U3-89S6C-C, Sony IMX255 sensor). The inverted dark-field microscope also housed a microscope objective (Olympus LUCPlanFL N 20x 0.45) to image the samples and piezo stages (Thorlabs PD2/M 5 mm Linear Stage with Piezoelectric Inertia Drive), allowing us to automate the system to systematically capture images of particles at higher throughput and with improved usability. The functionalised nanoparticles with complementary DNA handles were loaded into a custom-built coverslip microchamber formed using parafilm spacers. Using the automated imaging system, thousands of images were systematically captured across the chip. From now on, we will use NP to refer to both both single NP and clusters, as they are indistinguishable below the diffraction limit of our imaging system.

**Figure 2.**
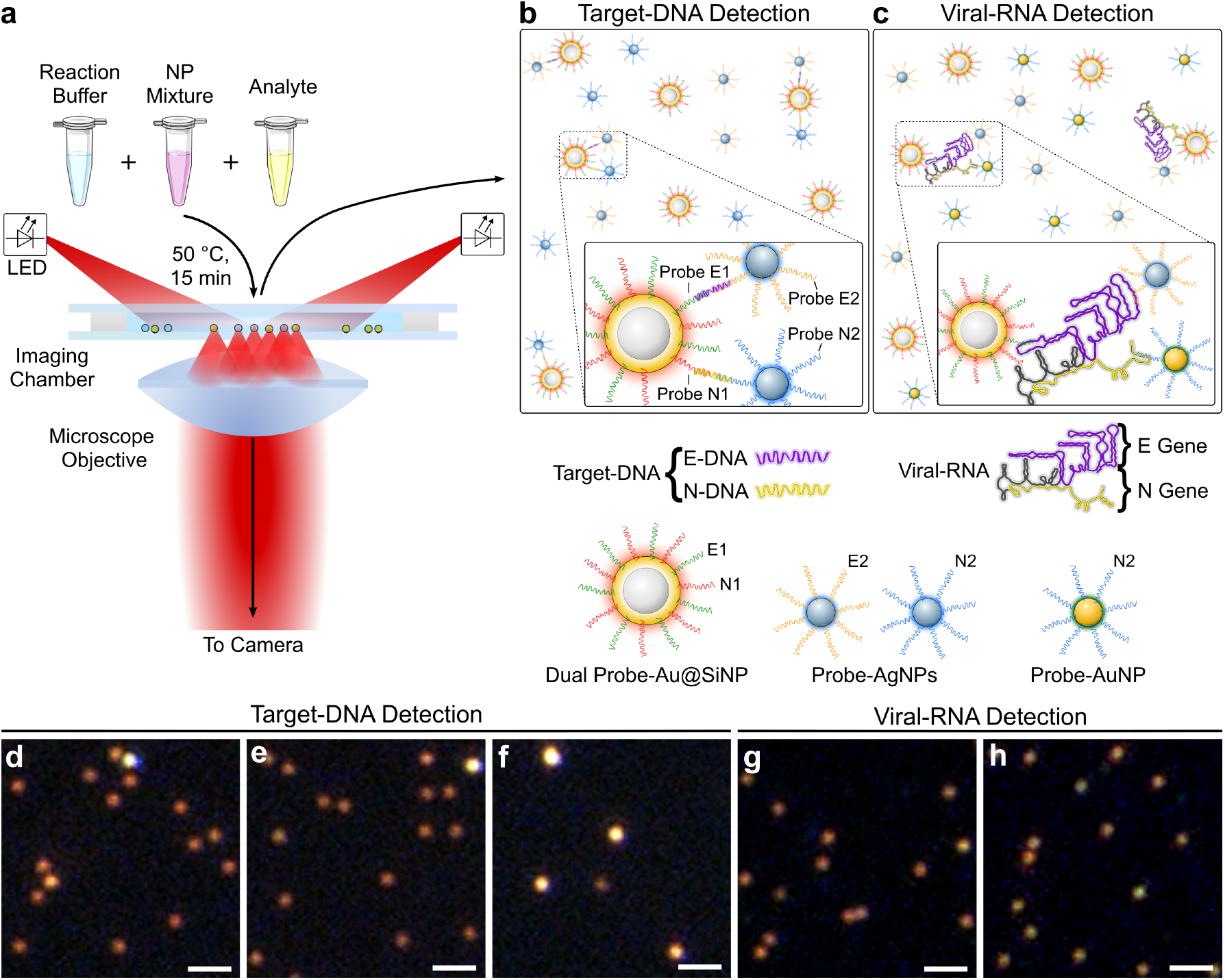
Nanoparticle imaging assay overview. (a) A schematic representation of the assay steps, with optically barcoded nanoparticles (NPs), analysed via dark-field microscopy. (b,c) Schematic representations of nanoprobes binding to Target-DNA (mixed ssDNA derived from SARS-CoV-2, see Supplementary Information Table S4) or Viral-RNA (genomic RNA from whole SARS-CoV-2), respectively. (d)-(f) Exemplary dark-field images (RGB) of the Target-DNA detection scheme, showing images captured for control (d), 500 fM Target-DNA ([E-DNA] = [N-DNA] = 500 fM) (e), and 50 pM Target-DNA ([E-DNA] = [N-DNA] = 50 pM) (f). (g,h) Exemplary dark-field images (RGB) of the Viral-RNA detection scheme, showing images captured for control (g) and 445 fM Viral-RNA (h). Scale bars = 5 µm.

To inform of our assay design, we exploited a combined equimolar mixture of a synthesized 188-bp single-stranded DNA (ssDNA) (“E-DNA”) derived from SARS-CoV-2 Envelope gene (*E* gene) and a 200-bp ssDNA (“N-DNA”) from SARS-CoV-2 Nucleocapsid gene (*N* gene), that we will henceforth refer to as “Target-DNA” ([Target-DNA] = [E-DNA] = [N-DNA]). By mixing the two ssDNA sequences, we attempted to approximate genomic RNA detection (where both genes are present). Despite obvious limitations, this choice allowed us to test the assay performance with well-characterized and stable target molecules before attempting to detect full-length SARS-CoV-2 RNA. It also simplified probe selection, as the same DNA probes could be used for both applications. For Target-DNA detection, we used Au@SiNPs and AgNPs only, with probes complementary to E-DNA and N-DNA (Figure 2b). The NPs were first mixed with a buffer formulated to reduce non-specific aggregation before adding Target-DNA (or blank buffer) at defined concentrations spanning an order of magnitude (50 fM, 500 fM, 5 pM, and 50 pM). Samples were then heated briefly (15 min) at 50 ^°^C before transfer to an imaging chamber for dark-field analysis. We utilize the beneficial characteristics of Au@SiNPs, which due to their larger mass will settle on the glass coverslip surface naturally by gravity. The Au@SiNP spots can then be easily localized and further analysed for their spectral characteristics to determine binding of AgNPs in a Target-DNA dependent manner (Figure 2d-f).

We next tested the assay with heat-inactivated SARS-CoV-2 virus (capsid intact, “Viral-RNA”) (Figure 2b). This introduced added complexity, as the virus was supplied in serum-fortified medium and required lysis. By incorporating high surfactant concentrations, we simultaneously stabilized NPs^28,29^ and lysed viral capsids^30^, hence enabling a one-pot format. To increase sensitivity, one AgNP probe was replaced with AuNPs to fully exploit the RGB colour space (Figure 2c). Released Viral-RNA could cross-link all three NP species, with Au@SiNPs acting as dual probes for intact genomes or *E*/*N* gene fragments. Virus dilutions of 111, 223, and 445 fM (genome equivalents) produced analyte-dependent clusters observable in dark-field images (Figure 2g,h). Detailed NP preparation, buffer composition, and imaging protocols are provided in the Methods section.

### 2.3 Statistical analysis

The spectral information of individual particles from the image datasets were analysed in Python. Using thresholding, we isolated NPs in each image and recorded their RGB intensity values. This analysis was performed on a representative subset of ≈600 images from the target-DNA assay, sampled across all concentrations and experimental days from a total dataset of ≈2500 images. In Figure 3, we plot the three colour channel intensities, finding that there is a clear correlation with the target-DNA concentration. However, the existence of more than one cluster for some concentrations suggests that minor variations in assay preparation and imaging conditions affect the NP agglomeration. These day-to-day experimental variations prevent us from doing robust classification, especially at low concentrations where it is difficult to separate the target-DNA and control. Therefore, we implemented a DL model to enable accurate and efficient concentration classification.

**Figure 3.**
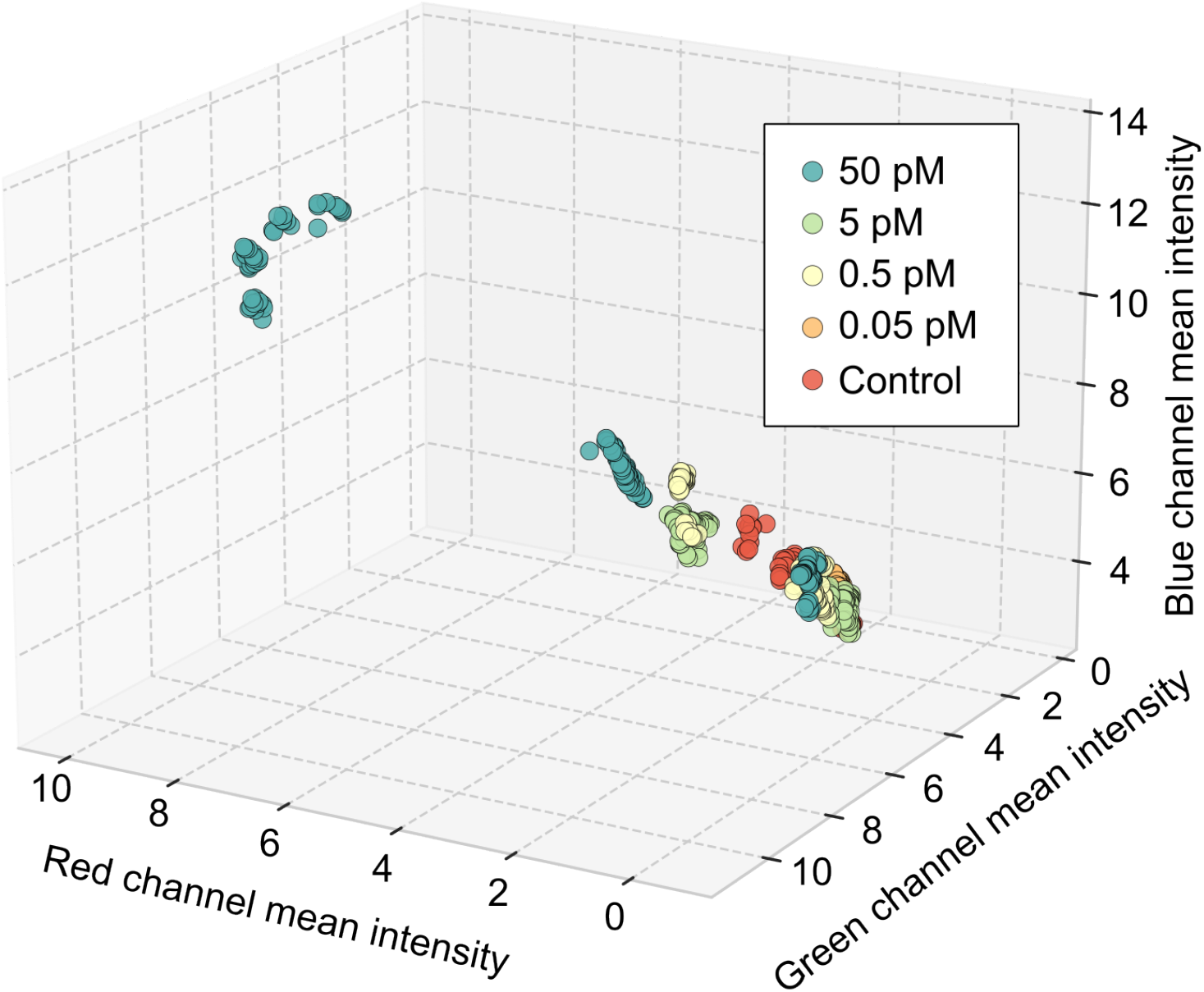
Three-dimensional scatter plot showing average red, green, and blue channel intensities, with each point representing particle averages within a single image.

### Deep Learning Workflow

To better classify biomolecular binding events using subtle spectral variations of NPs, we use a supervised DL approach, which automates the counting and differentiation of NP sub-populations. To effectively capture spectral and morphological information in a NP-based microscopy assay, we propose a new Mc-GNN architecture. For concentration prediction, integer labels encoded each analyte concentration. Raw datasets were first processed as shown in Figure 4, where each dark-field image was preprocessed to isolate single NPs through a global background correction, CLAHE-based contrast enhancement^31^, and Gaussian blur for contour detection. Particle-wise background correction was then applied by estimating local background in an expanded region around each particle, while masking out nearby particles. Radial sorting of contours from the optical centre outward prioritized particles least affected by defocussing effects and vignetting^32,33^. The sorting across raw images also facilitates the incorporation of spatial information into the analysis. These steps compensated for illumination drift and detector offsets, which often occur in realistic experimental environments, producing uniformly focused, intensity-balanced crops without explicit need for an ROI alignment stage. Each segmented particle was resized to [3, 15, 15], and all particles from a raw image (*N ≈* 1, 000-5, 000) were stacked into batched input tensors of shape [*N*, 3, 15, 15].

**Figure 4.**
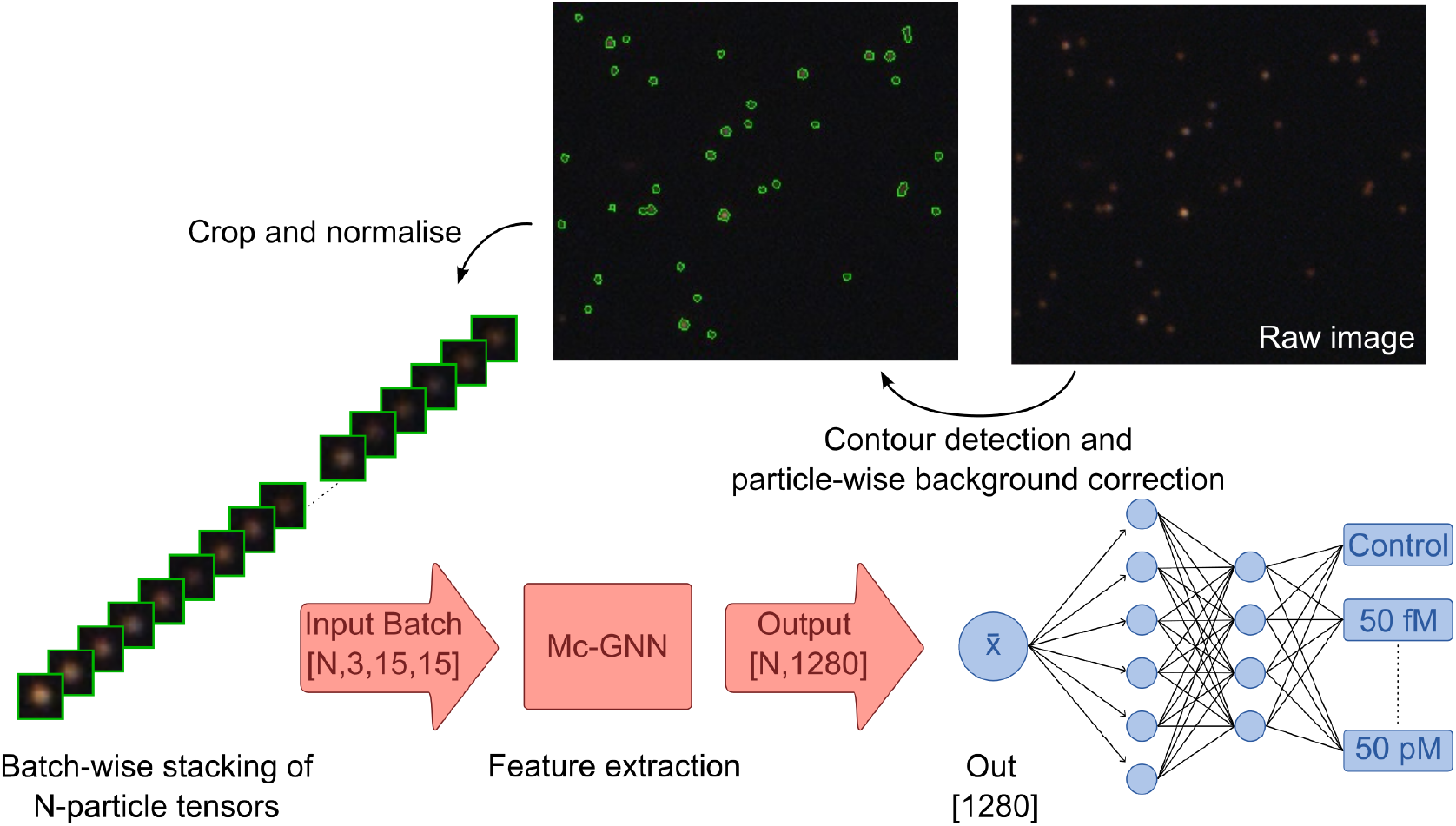
DL analysis pipeline. Raw images undergo global correction to support contour detection (steps not depicted in this figure), followed by particle-wise background correction, radial sorting, and cropping. All the N-particle tensors are stacked in shape [*N*, 3, 15, 15] and processed by Mc-GNN for feature extraction, followed by fully connected layers for the concentration prediction.

The resulting particle tensors were inputs into the Mc-GNN model, whose architecture is shown in Figure 5. The model consists of two main components: (i) a morphology-guided convolutional stage that uses five structural masks that include circular, Gaussian, doughnut, square, and edge-for morphology-aware feature extraction; and (ii) a graph-based relational modelling that models morphology-guided kernel outputs from step (i) as a Graph, which is then followed by the GCN layer (Graph Convolutional Network; a common layer in GNNs that aggregates neighbouring features). This results in relational learning of meaningful interactions between multiple morphology-related features at the particle level through a graph-based feature propagation. Each of these masks emphasize distinct structural properties relevant to the NP morphology: **Circular (C)**: emphasizes isotropic particle intensity, **Gaussian (G)**: captures centre-focused (diffuse or gradient-like structures), smooth intensity distributions, **Doughnut (D)**: highlights ring-like particle structures, **Square (S)**: captures rectilinear or grid-like features, **Edge (E)**: enhances particle boundaries. Unlike many DL models, our Mc-GNN is therefore a relatively intuitive model; every structural mask highlights a distinct attribute that the model extracts from an individual particle and, following graph-based steps, does relational learning to enhance the extracted information. The resulting particle-level embeddings were averaged on an image-level and subsequently passed through a fully connected layer for concentration prediction. Training was performed with the dataset split of 80% training/validation(3:1) and 20% testing in PyTorch 2.4.1 with CUDA 12.1 on an NVIDIA RTX A5000 GPU, for 300 epochs, using the Adam optimizer (1 × 10^−5^ learning rate) with early stopping (patience of 50).

**Figure 5.**
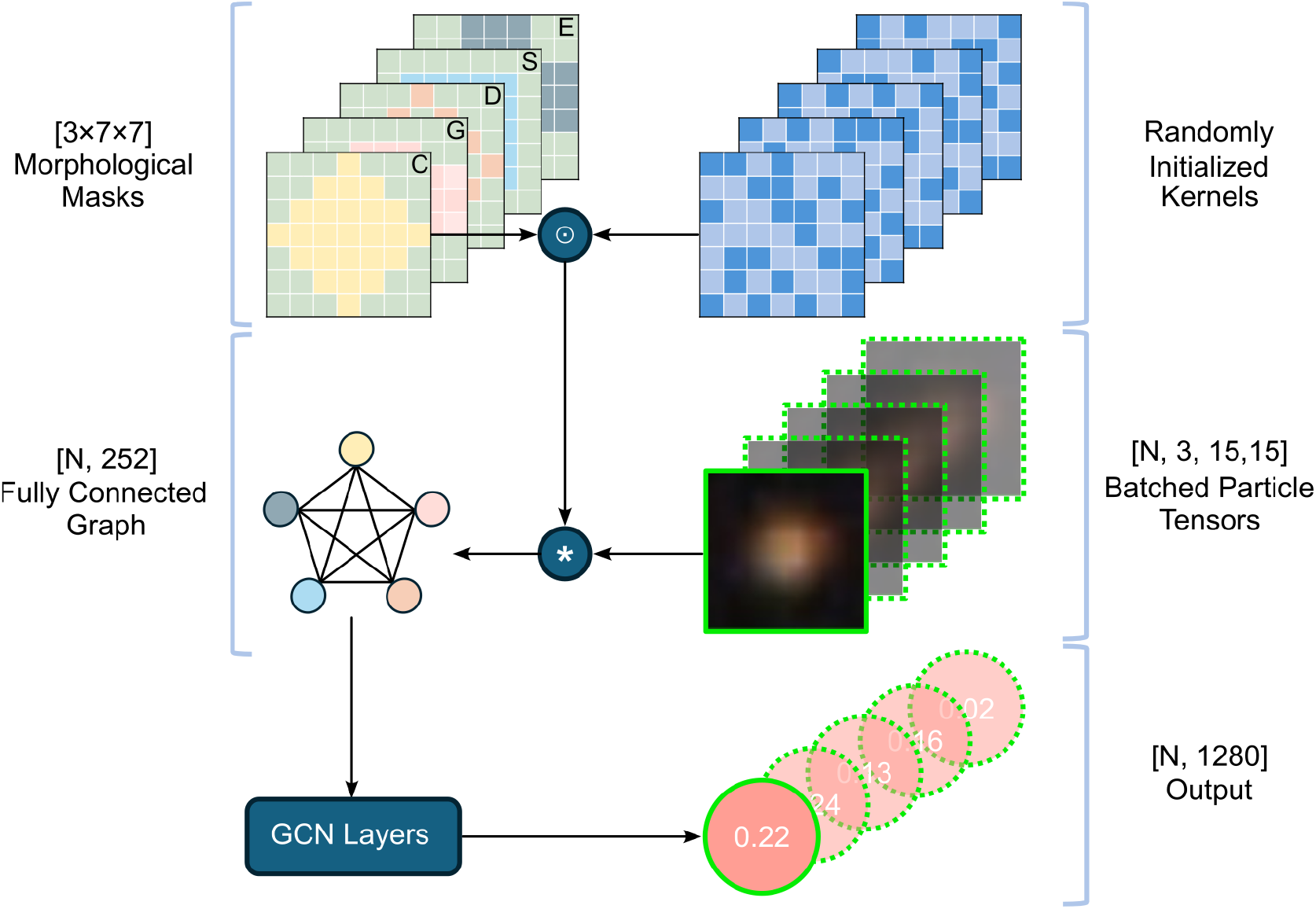
Architecture of the Mc-GNN model. Each input raw image is preprocessed into batched particle tensors [*N*, 3, 15, 15], which are convolved with randomly initialized kernels modulated by defined morphological masks - C: Circular, G: Gaussian, D: Donut, S: Square, E: Edge, 252 of each sized [3 × 7 × 7]. Spatially compressed resulting feature embeddings [*N* × 5, 252] form nodes in a fully connected graph, where graph-based relational learning across morphology-guided kernel outputs is performed via the GCN layers. This yields final particle-wise outputs [*N*, 1280] for downstream predictions. “⊙” denotes Hadamard (element-wise) product; “*” denotes convolution.

### 2.4 Classification based on deep learning model

Model performance was assessed using Precision, macro Recall (referred to as “Recall”, same as average class-wise accuracy), and macro F1-score (referred to as “F1-score”) averaged across 5-fold stratified cross-validation. Area under the ROC curve (AUC) was excluded here due to the multiclass structure and class imbalance. Model comparisons used Friedman tests (as indicated by the *χ*^2^ statistic and *p*-value), which was followed by Wilcoxon signed-rank post-hoc tests to identify which model pairs showed statistically significant differences^34^. All models were trained and validated using 5-fold stratified cross-validation with consistent preprocessing and data partitions, as described previously. Although we did not test additional random seeds, the consistent cross-validation results across folds confirms that the model’s performance was stable, as reflected by Friedman rankings of 1.0 across subsequent ablation studies, and comparative performances.

As displayed in Figure 6, in the cross-validation fold with the highest performance (note that the average across all folds was later used for comparative analyses), the Mc-GNN achieves a high recall, where it accurately classifies all images of 50 fM and 500 fM, but also 98.4% of control images, 98.9% of 5 pM images, and 99.2% of 50 pM images (Figure 6a). On the Viral-RNA dataset, the model maintained high accuracy, correctly classifying 111 fM samples while achieving 91-96% recall across the other concentrations. Additional clinically relevant binary analyte detection results are reported in the SI, Fig. S1. These results highlight that the main advantage of the DL model is its ability to efficiently and accurately classify the SARS-CoV-2 concentration based on our NP assay, even when the feature-level information is difficult to analyse with conventional analytical methods.

**Figure 6.**
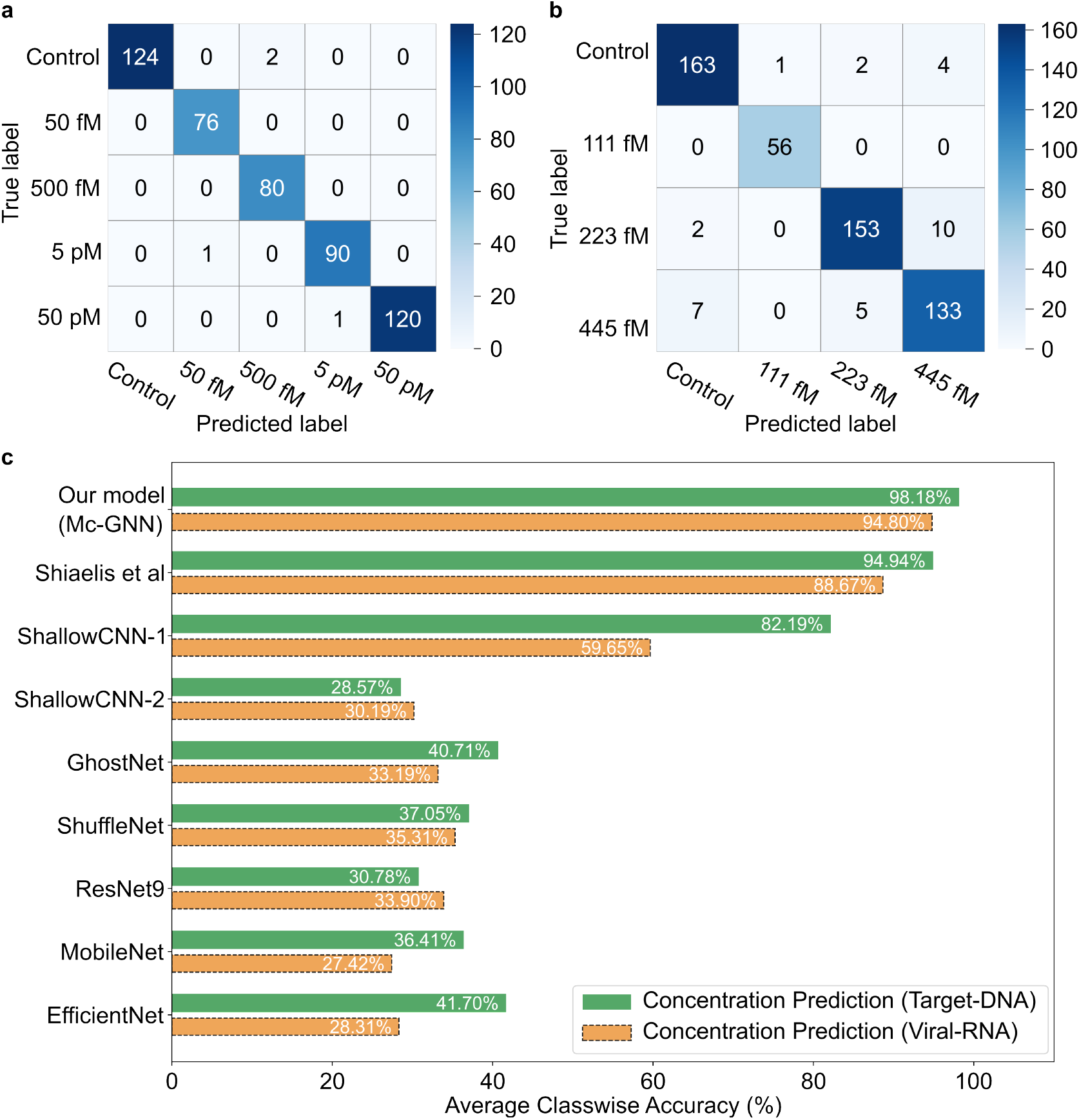
Results of cross-validation fold with the highest performance: (a) Confusion matrix for Target-DNA concentration prediction. (b) Confusion matrix for Viral-RNA concentration prediction. (c) Comparative performance of different models on both datasets.

We benchmarked Mc-GNN against other DL architectures (ShuffleNet^35^, MobileNet^36^, GhostNet^37^, EfficientNet^24,38^, ResNet9^39^(higher variants as in McAffee *et al*.^25^ were computationally prohibitive in our setting), and domain-specific DL baselines. Figure 6c and Tables 1 and 2 summarize the comparative performance of all evaluated models on the Target-DNA and Viral-RNA concentration prediction tasks. The Friedman tests revealed significant performance differences across models for all evaluated metrics (precision, recall, F1-score; p<10^−4^), with Mc-GNN consistently achieving the top rank (1.0) across both datasets. Specifically, Mc-GNN achieved 98.2% average class-wise recall accuracy on the Target-DNA dataset and 94.8% on the Viral-RNA dataset, with statistically significant improvements over all baselines confirmed by post-hoc Wilcoxon signed-rank tests (p=0.0312). The fluorescence-based model of Shiaelis *et al*.^21^ ranked second, achieving 94.9% and 88.7% accuracy, respectively, demonstrating the competitive power of particle-resolved fluorescence models, but with the assistance of fluorescence labels required. Most other baselines performed substantially worse, below 83% for Target-DNA and 60% for Viral-RNA. ShallowCNN-1 (by Sajed *et al*.^11^) for example achieved moderate results, confirming that 2D convolutional architectures are unable to fully capture complex clustering patterns. This limitation was further evidenced by the reduced performance in ShallowCNN-2^10,22,23^, where an increasing the number of convolutional layers led to worse performance. More compact architectures (ResNet9, GhostNet, ShuffleNet, MobileNet, EfficientNet) showed unstable precision–recall trade-offs, leading to markedly lower F1-scores (≤ 0.35 for DNA and ≤ 0.20 for RNA tasks), indicating that these DL models were unsuitable for these types of datasets. We excluded the AlexNet variant as proposed by Wang *et al*.^20^ and the adaptive FPN model^19^, as they were designed for full-field microscopy and incurred prohibitive memory costs in our particle-level setting. See the SI (Section S1) for detailed discussions of Tables 1 and 2. Unlike conventional image-level models, Mc-GNN intuitively integrates the important nanoparticle attributes, such as their spectral, spatial (distribution/spacing), and morphological (size/shape) properties, by explicitly integrating morphology-guided kernel convolution with relational graph modelling. A brief exploration of alternative hyperparameters (such as kernel size and the number of kernels) yielded no meaningful improvements to our model for the analyte detection task, so we did not perform further extensive tuning. Related works on morphology-guided convolutional kernels^40–42^ primarily target computationally intensive semantic segmentation pipelines^43–51^. While, Graph-based approaches such as KerGNN^52^, dual-graph CNNs^53,54^, and hybrid CNN-GNN frameworks (e.g., SR-GNN, CGC-Net, MorphoGNN^55–58^), as well as classic GNNs^59–61^, lack explicit morphology-aware constraints at the initial convolution stage. Overall, these findings demonstrate that morphology-aware convolution with graph modeling is a critical factor for accurate and efficient analyte concentration prediction in nanoparticle-based diagnostic assays, all while using under 5 GB GPU memory versus > 10 GB for most architectures for up to 5,000 particles per image.

**Table 1.** Performance comparison of the Mc-GNN with contemporary models on the Target-DNA dataset for the concentration prediction task. Friedman rankings and Wilcoxon signed-rank test *p*-values (vs. Mc-GNN) are reported.

| Model | Precision |  |  | Recall |  |  | F1-Score |  |  |
| --- | --- | --- | --- | --- | --- | --- | --- | --- | --- |
|  | Score | Rank | *p*-val | Score | Rank | *p*-val | Score | Rank | *p*-val |
| **Mc-GNN** | **0.9813** | **1.0** | - | **0.9818** | **1.0** | - | **0.9814** | **1.0** | - |
| Shiaelis *et al.*<sup>21</sup> | 0.9484 | 2.0 | 0.0312 | 0.9494 | 2.0 | 0.0312 | 0.9475 | 2.0 | 0.0312 |
| ShallowCNN-1<sup>11</sup> | 0.8315 | 3.0 | 0.0312 | 0.8219 | 3.0 | 0.0312 | 0.8170 | 3.0 | 0.0312 |
| ShallowCNN-2<sup>10,22,23</sup> | 0.4104 | 6.8 | 0.0312 | 0.2857 | 8.0 | 0.0312 | 0.2104 | 8.0 | 0.0312 |
| GhostNet<sup>37</sup> | 0.5183 | 6.2 | 0.0312 | 0.4071 | 5.6 | 0.0312 | 0.3493 | 5.8 | 0.0312 |
| ShuffleNet<sup>35</sup> | 0.4768 | 7.2 | 0.0312 | 0.3705 | 5.8 | 0.0312 | 0.3356 | 5.0 | 0.0312 |
| ResNet9<sup>39</sup> | 0.6081 | 5.2 | 0.0312 | 0.3078 | 7.8 | 0.0312 | 0.1884 | 8.0 | 0.0312 |
| MobileNet<sup>36</sup> | 0.3979 | 7.8 | 0.0312 | 0.3641 | 6.6 | 0.0312 | 0.2948 | 6.4 | 0.0312 |
| EfficientNet<sup>24,38</sup> | 0.5298 | 5.8 | 0.0312 | 0.4170 | 5.2 | 0.0312 | 0.3482 | 5.8 | 0.0312 |

**Table 2.** Performance comparison of the Mc-GNN with contemporary models on the Viral-RNA dataset for the concentration prediction task.

| Model | Precision |  |  | Recall |  |  | F1-Score |  |  |
| --- | --- | --- | --- | --- | --- | --- | --- | --- | --- |
|  | Score | Rank | *p*-val | Score | Rank | *p*-val | Score | Rank | *p*-val |
| **Mc-GNN** | **0.9476** | **1.0** | - | **0.9480** | **1.0** | - | **0.9470** | **1.0** | - |
| Shiaelis *et al.*<sup>21</sup> | 0.8661 | 2.0 | 0.0312 | 0.8867 | 2.0 | 0.0312 | 0.8728 | 2.0 | 0.0312 |
| ShallowCNN-1<sup>11</sup> | 0.5939 | 3.6 | 0.0312 | 0.5965 | 3.0 | 0.0312 | 0.5811 | 3.0 | 0.0312 |
| ShallowCNN-2<sup>10,22,23</sup> | 0.4085 | 6.8 | 0.0312 | 0.3019 | 6.6 | 0.0312 | 0.2224 | 6.4 | 0.0312 |
| GhostNet<sup>37</sup> | 0.4302 | 7.2 | 0.0312 | 0.3319 | 6.2 | 0.0312 | 0.2073 | 6.8 | 0.0312 |
| ShuffleNet<sup>35</sup> | 0.3784 | 7.2 | 0.0312 | 0.3531 | 5.4 | 0.0312 | 0.3071 | 4.0 | 0.0312 |
| ResNet9<sup>39</sup> | 0.4776 | 5.6 | 0.0312 | 0.3390 | 5.8 | 0.0312 | 0.1946 | 7.6 | 0.0312 |
| MobileNet<sup>36</sup> | 0.5319 | 5.0 | 0.0312 | 0.2742 | 7.6 | 0.0312 | 0.1708 | 7.6 | 0.0312 |
| EfficientNet<sup>24,38</sup> | 0.4361 | 6.6 | 0.0312 | 0.2831 | 7.4 | 0.0312 | 0.1957 | 6.6 | 0.0312 |

### Ablation Study

To evaluate the contribution of individual architectural components in our Mc-GNN model, we conducted an ablation study on the concentration prediction task using both the Target-DNA and Viral-RNA datasets. The baseline variants included: *(i)* Morphology-Guided Convolution, *(ii)* Fixed-Shape Kernel Convolution, and *(iii)* Fixed-Shape Kernels with GNN. A detailed ablation table with the detailed results of significance tests is provided in the SI (Table S3). We find that the Mc-GNN achieved the best average rank (1.0) across all metrics (F1 = 0.9814 for Target-DNA dataset) and in comparison, removal of the morphology-guided convolution resulted in performance drops to F1 = 0.9103 and F1 = 0.8729 for Fixed-Shape Kernels + GNN and for Fixed-Shape Kernel Convolution, respectively. Similar reductions are seen for Viral-RNA dataset (see Table S3). This shows model variants that either remove morphology guidance or defer it to later stages (Fixed-Shape Kernels + GNN, similar to generic CNN+GNN stacks) underperform, indicating that post hoc graph aggregation alone does not perform better. A morphology-only variant (Morphology-Guided Convolution without GNN) shows an improved performance (F1 = 0.9643) but still trails the full model, highlighting the benefits of relational reasoning. Statistical tests confirm significant differences among variants, emphasizing the interdependence of morphology-guided kernels and graph-based feature aggregation for classifying NP aggregation assays.

## Conclusions

Colorimetric nanoparticle assays offer a promising solution toward developing a diagnostic tool that is simultaneously sensitive, cost-effective, user-friendly and chemically simple. So far, no existing diagnostic test has been able to achieve all of these attributes. In this work, we demonstrate a promising route towards this goal, by combining nanoparticles with distinct scattering characteristics, low-magnification optics and a deep learning algorithm for the image analysis. Although image-based approaches using consumer optics and machine learning have previously shown promising results, most rely on complex assay protocols or, on the analysis side, on whole-image classification. They often lack particle-level analysis without external labelling and incur heavy computational resources, hence hindering their practical implementation^19–22^. Our calculations show that distinct nanoparticles and their interparticle distance–dependent LSPR shifts make it possible to distinguish specific from non-specific binding events. We deliberately maintain a simple, one-pot, fluorescence-, amplification- and enzyme-free assay design^7–9,62–64^ that can be loaded on to a small chamber on a custom coverslip chip, while capturing the spectral information with a standard RGB camera. Although particle-level analysis reveals a correlation between SARS-CoV-2 DNA concentration and the red, green and blue channel mean intensity, day-to-day experimental variability renders this feature unreliable for robust concentration classification. To address these realistic experimental conditions, we develop a Mc-GNN architecture that performs particle-level analysis with context-aware relational learning and feature aggregation. By explicitly including morphological sensitivity into our model, we are able to outperform similar DL models when it comes to NP image analysis and we show that our platform can perform femtomolar DNA and RNA detection. Mc-GNN is able to resist imaging noise, day-to-day experimental variations and optical aberrations, and its design enables straightforward extension to additional analytes, NP chemistries, or multiplexed modalities. All of this while remaining computationally efficient on consumer-grade GPUs (< 5 GB RAM).

Looking ahead, the demonstrated cross-task and cross-modality robustness of Mc-GNN highlights its promise for even broader diagnostic applications. Examples include its extension to more diverse datasets and an increased input concentration granularity will allow the Mc-GNN to be applied as a regression-based model for continuous concentration quantification. Clinical validation on real patient-derived samples, including blood or saliva, will also be critical for translation and benchmarking against standard assays, such as LFAs and RT-PCRs. Although currently less sensitive than RT-PCR (<100 copies/mL^65^), our purification-free assay is expected to improve with additional data and avoids the need for rigorous sample purification^26,27^. Finally, further miniaturisation of our assay using a portable imaging device combined with an on-device AI application could democratize molecular diagnostics by providing high-performance testing to decentralized or resource-limited environments.

## Methods

### DNA Functionalization of Gold and Silver Nanoparticles

Spherical gold nanoparticles (AuNPs) (diameter, 50 nm), spherical silver nanoparticles (AgNPs) (diameter, 50 nm), and gold-shell silica-core nanoparticles (Au@SiNPs) (80 nm Si core, 20 nm Au shell) each with 40 kDa polyvinylpyrrolidone (PVP) as capping agent were purchased from nanoComposix (USA) (Figure S2). NPs were functionalized with thiolated DNA probes (see SI Table S4) (Integrated DNA Technologies, USA) using a modified PVP-assisted conjugation protocol^66,67^. Probe DNA sequences were chosen semi-empirically for particle stability, with initial designs created using NUPACK Web Application^68,69^ and with subsequent testing at high salt after conjugation. Thiolated DNA was reduced, for use with AgNPs only, by incubation with tris(2-carboxyethyl)phosphine hydrochloride (TCEP) (Sigma Aldrich) at a 1:100 DNA:TCEP ratio for 1 hr at room temperature. DNA for use with gold was not treated, as TCEP reduction was found to be unnecessary^70^. For conjugation, PVP-capped NPs were first aliquoted from high concentration stocks (AuNP: 27.9 OD @ 528 nm, *est*. 1.2 nM; AgNP: 134.9 OD @ 419 nm, *est*. 2.7 nM; and Au@SiNP: 37.8 OD @ 666 nm, *est*. 0.15 nM) into DNA Lobind tubes (Eppendorf SE, Germany). The respective thiolated probes were added at a final concentration of 28.5 µM followed by bath sonication of the reaction (10 s)^29^ and incubation (10 min) at room temperature. Additional PVP (40 kDa) (Sigma Aldrich) was added to a final concentration of 0.35% (w/v) to further stabilize the particles. A volume of salt-containing buffer, consisting of 20 mM sodium phosphate buffer (PB, pH 8), 200 mM NaBr, and 0. 02% (w/v) sodium dodecyl sulfate (SDS) (Sigma Aldrich), equivalent to the reaction volume was slowly added in a single step. Notably, NaBr was used instead of NaCl to discourage nonspecific adhesion of nucleotides to the gold/silver surface^71^. The reaction mixture was then heated at 60 °C for 2.5 hr with mixing (800 rpm). To remove residual DNA and salts, the resulting conjugated NPs were washed 5 times (or 7 times for viral detection) by centrifugation (18000*g*, 5 min) using 10 mM PB (no NaCl). Conjugated NP concentrations were then approximated using their UV-Vis Absorbance spectra (NanoDrop One Instrument, Thermofisher Scientific, USA) as compared to their reported stock concentrations.

### DNA detection

A 188 bp single-stranded DNA sequence (“E-DNA”) derived from the Envelope Gene and 200 bp DNA sequence (“N-DNA”) derived from the Nucleocapsid Gene of SARS-CoV-2 (NCBI NC_045512.2) were designed for use as model DNA targets (see SI Table S4) (Ultramer ssDNA, PAGE purified; Integrated DNA Technologies, USA). We used these two strands at equivalent concentrations to emulate a larger SARS-CoV-2 target and refer to them collectively as “Target-DNA.” For Target-DNA detection, dual-probe Au@SiNPs (PrE1/PrN1-Au@SiNP) as well as two single-probe AgNP species (PrE2-AgNP, PrN2-AgNP) were used for simultaneous detection of E-DNA and N-DNA (Table S4). First, a NP mixture (in PB), a reaction buffer mixture, and Target-DNA (in TE buffer) at the relevant concentrations were prepared. The three separate solutions were mixed to start the hybridization assay. The reaction was set up such that the following final concentrations were achieved in a 40 µL volume: 10 mM PB, 1 pM Au@SiNP-PrE1, 3 pM AgNP-PrE2, 3 pM AgNP-PrN2, 400 mM NaCl, 2 mM Ethylenediaminetetraacetic acid (EDTA) (Sigma Aldrich), 3% (w/v) dextran (150 kDa) (Sigma Aldrich), 0.05% (w/v) PVP (40 kDa), 0.05% (w/v) SDS, and lastly 50 fM - 50 pM Target-DNA ([E-DNA] = [N-DNA] = reported concentration) or TE buffer as control. EDTA was included to chelate undesirable multivalent ions, while dextran was included to promote specific hybridization^72,73^. Salt and surfactant concentrations were empirically derived for optimal hybridization results (fast and specific). Immediately after mixing, the solution was vortexed and then heated to 50 °C with mixing (800 rpm) for 15 min. The reaction was allowed to cool for 10 min at room temperature before transferring 20 µL to a home-made microscopy chamber consisting of two plasma-etched coverslips sandwiching a PDMS well for optical imaging. Particles were imaged in solution using the optical setup detailed below. Notably, single Au@SiNPs settle via gravity onto the glass surface, while other particles tend to stay diffuse in solution unless they are clustered. Images were captured in an automated fashion using a piezo stage (see below) in order to generate sufficient training data for DL analysis.

### Viral RNA detection

For viral RNA detection, the following reagent was deposited by the Centers for Disease Control and Prevention and obtained through BEI Resources, NIAID, NIH: SARS-Related Coronavirus 2, Isolate USA-WA1/2020, Heat Inactivated, NR-52286. Viral load was determined by BEI Resources using droplet digital PCR (stock concentration: 5.36 × 10^8^ genome equivalents / mL). Virus was used either directly as provided or diluted in its storage medium, which consisted of Eagle’s Minimal Essential Medium (EMEM) (ATCC, USA) with 2% (v/v) fetal bovine serum (FBS) (Thermo Fisher Scientific, USA). Blank viral storage medium was used for virus dilution and as a control. For this assay iteration, dual-probe Au@SiNPs (PrE1/PrN1-Au@SiNP), single-probe AgNPs (PrE2-AgNP), and single-probe AuNPs (PrN2-AuNP) were used (Table S4). Therefore, the Viral-RNA could potentially cross-link all three particle species. First, a NP mixture (in PB), a reaction (lysis) buffer mixture, and the target/control (in viral storage medium) at the relevant concentrations were prepared. The three separate solutions were mixed to start the viral detection assay. The reaction was set up such that the following final concentrations were achieved in a 20 µL volume: 10 mM PB, 1 pM Au@SiNP-PrE1/PrN1, 3 pM AgNP-PrE2, 3 pM AuNP-PrN2, 200 mM NaCl, 50 mM EDTA, 3% (w/v) dextran sulfate (500 kDa) (Sigma Aldrich), 0.5% (w/v) SDS, 0.5% Triton-X100 (Sigma Aldrich), 10 mg/mL Proteinase K (Thermo Fisher Scientific, USA) and lastly 10 µL of inactivated SARS-CoV-2 at the relevant dilution (or viral storage medium as control). SDS, Triton-X100, and Proteinase K were included at relevant concentrations for viral lysis and nuclease inhibition^74^. Upon mixing, the sample was treated as described above for heating and imaging.

### Optical imaging

Images, including spectral information, of different NP sub-populations was collected using a colour (RGB) camera (Teledyne FLIR Grasshopper 3, model GS3-U3-89S6C-C, Sony IMX255 sensor). The camera was integrated into a custom-built inverted dark-field microscope that also housed broad-spectrum light-emitting-diodes (LEDs) (Cree C503D-WAN, 40k mcd intensity) as an affordable light source and a microscope objective (Olympus LUCPlanFL N 20x 0.45) to image the samples. Subtle spectral variations are classified using a supervised DL approach, which automates the counting and differentiation of NP sub-populations. To generate labeled training data for the DL model, we implemented an imaging protocol to systematically capture images of particles in an automated fashion using piezo stages (Thorlabs PD2/M 5 mm Linear Stage with Piezoelectric Inertia Drive).

### Proposed architecture and Implementation

To accurately capture particle-level spectral, spatial and morphological information in NP-based assays for concentration prediction, we propose Mc-GNN, a hybrid convolutional-graph architecture (Figure 5).

#### Input and Morphology-guided Kernel Design

Following the input raw image preprocessing and the dataset preparation steps, the input image is processed as batched tensor **X** ∈ ℝ^*N*×3×15×15^, where *N* denotes the number of particle images in a batch. To embed domain-specific inductive biases into the learning process, we introduce a set of shape-aware masked convolutional kernels. Specifically, five structural masks-*circular, edge, Gaussian, donut, and square*-are utilized to modulate the learnable kernels. Each of these masks emphasizes distinct structural properties relevant to NP morphology: **Circular (C)**: emphasizes isotropic particle intensity, **Gaussian (G)**: captures center-focused (diffuse or gradient-like structures), smooth intensity distributions, **Donut (D)**: highlights ring-like particle structures, **Square (S)**: captures rectilinear or grid-like features, **Edge (E)**: enhances particle boundaries. Randomly initialized 252 kernels **K** ∈ ℝ^252×3×7×7^ are element-wise modulated by 252 of each of these 5 masks **M**^(*s*)^, producing morphology-guided kernels. The selection of 252 kernels per shape was empirically determined via non-extensive hyperparameter tuning to optimize model performance. After normalization, convolution of these morphology-guided kernels with input particle tensors directly encodes morphological priors into the resulting feature maps, making them explicitly shape-aware at the earliest feature extraction stage. Spatial global average pooling compresses feature maps into compact vectors. The pooled outputs from all five shape branches to form are stacked to form a unified tensor of shape [*N*, 5, 252], which is then permuted into the shape [*N* *5, 252] to facilitate downstream graph-based relational modelling.

#### Graph-based relational modelling and GCN based relational learning

The spatially compressed and permuted tensor 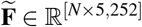 encodes, for each particle, five parallel morphology-guided features, each as a 252-dimensional embedding derived from a distinct morphology-guided kernel branch. This multi-view representation serves as a foundation for capturing context-aware relational learning across morphology-guided kernel outputs. Thus, corresponding to each input particle, we represent the above tensor as a graph with 5 nodes, where each is a 252-dimensional morphology-guided embedding vector. The inter-relations between these morphology-guided kernel outputs for each particle are captured via a learnable adjacency matrix **A** ∈ R^5×5^, where each entry *A*_*i j*_ denotes the importance of information flow from node *j* to node *i*. **H**^(*ℓ*+1)^ = *σ* (**ÂH**^(*ℓ*)^**W**^(*ℓ*)^), where **Â** is a normalized adjacency matrix learned to represent kernel interactions, **W**^(*ℓ*)^ are learnable weights, and *σ* (·) denotes a ReLU activation.

Two GCN layers act on the graph representation of morphology-guided particle-feature embeddings to produce final particle embeddings **z** ∈ ℝ ^[252*5=1280]^ upon flattening node representations. A GCN layer aggregates and transforms features from a node’s neighbors in a graph, enabling context-aware relational reasoning by capturing structural dependencies and shared context across connected nodes. Thus, this graph-based relational modeling passed through GCN layers enables Mc-GNN to dynamically capture complementary interactions among morphology-guided kernel outputs. By leveraging the unique strengths of each kernel type through GCN-based relational learning, the architecture adapts to diverse binding patterns and concentration conditions-enhancing interpretability and discriminative power in concentration prediction for the NP-based assays.

### Deep Learning Evaluation Protocol

Model performance was quantitatively assessed using standard classification metrics, including Precision, macro Recall (Recall), and F1-score. Macro Recall is computed by first calculating the recall for each class independently, and then averaging these values across all classes. This treats each class equally, regardless of its frequency in the dataset, and is therefore particularly appropriate for imbalanced classification problems. In the context of our concentration prediction task, macro Recall is equivalent to average class-wise accuracy, as it reflects the model’s ability to correctly identify each concentration class on a per-class basis before averaging. F1-score, the harmonic mean of Precision and Recall, was also computed for each class and then averaged (macro F1-score) to provide a balanced measure of performance that accounts for both false positives and false negatives. This is particularly useful when evaluating multiclass classification under class imbalance, where precision-recall tradeoffs may vary across classes. For all metrics, scores were computed on the held-out test set for each fold in a 5-fold stratified cross-validation scheme, and the final reported values represent the mean performance aggregated across all folds. Due to the multiclass and imbalanced nature of the concentration prediction task, AUC was not reported for this setting.

To assess the statistical significance of observed performance differences among models, we employed a rigorous non-parametric evaluation protocol. The Friedman test was first applied to the performance scores of all models across the cross-validation folds, with ranks computed to summarize each model’s relative performance. The Friedman test evaluates whether the observed ranking differences across models are likely to have occurred by chance; statistical significance is established if the test yields a low *p*-value (e.g., *p* < 0.05) and a large *χ*^2^ statistic, indicating that at least some models perform differently from others. Upon detecting significant overall differences, we conducted Wilcoxon signed-rank post-hoc tests^34^ for robust pairwise comparison between the proposed Mc-GNN and each baseline. These tests were performed on the fold-wise metric distributions obtained from identical data splits for all models. This non-parametric framework not only mitigates the impact of outliers and non-Gaussian metric distributions, but also minimizes bias arising from sampling variation, thereby providing a statistically robust foundation for comparative claims.

## Author contributions statement

K.J., X-L.C., and J.B.D. contributed equally to this work. K.J., X-L.C., J.B.D., P.T. and L.W. performed conceptualization. X-L.C., J.B.D., L.A.P., S.B., and F.N.N.M.H. performed experimental investigation. K.J. developed the data processing and deep learning method, carried out model training, and analyzed the results. Y.Y. and Y.W. contributed to model-related discussions. P.T. and L.W. performed supervision. K.J., X-L.C., and J.B.D. wrote the original manuscript draft. All authors contributed to review and editing of the final manuscript.

## Additional information

**Competing interests** The author(s) declare no competing interests.

**Data availability** The datasets generated during and/or analysed during the current study are available from the corresponding author on reasonable request.

**Code availability** The codes generated during the current study are available from the corresponding author on reasonable request.

