## Supplemental information for "Morphology-Guided Deep Learning for Nanoparticle Agglomeration Diagnostic Assays"

#### This PDF file includes:

Supporting text
Figs. S1 to S2
Tables S1 to S4
SI References

### Supporting Information Text

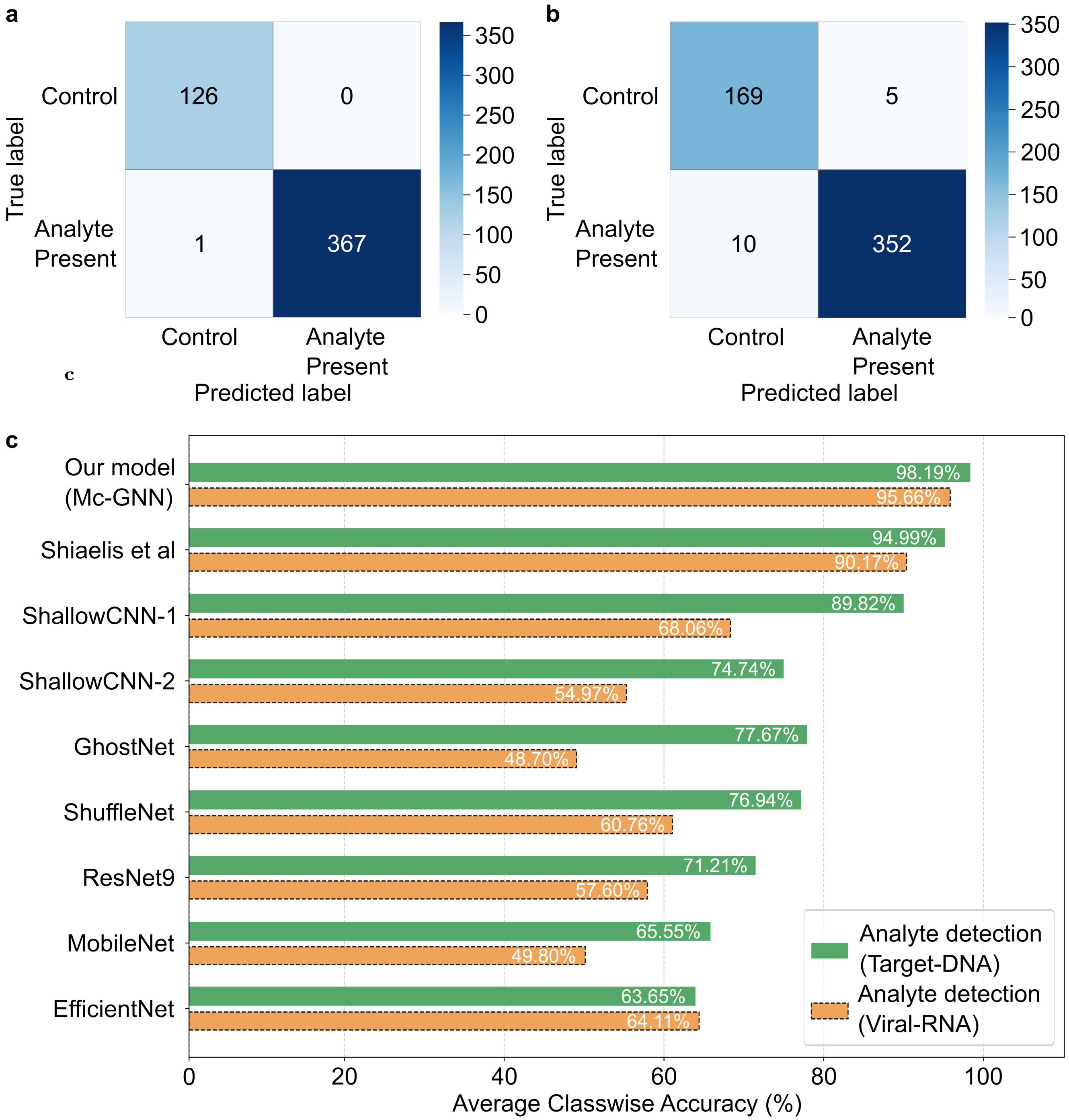


**Fig. S1.** Best fold test results: (a) Confusion matrix for Analyte Detection on Target DNA dataset. (b) Confusion matrix for Analyte Detection on Viral RNA dataset. (c) Comparative performance of Analyte Detection on different models.

### Additional Results

**Analyte Detection Performance Analysis .** In addition to concentration prediction, we first evaluated all models on the clinically relevant analyte detection task, serving as a foundational assessment of diagnostic efficacy. In this, the control class (absence of target analyte) was compared against all non-control concentrations, which were collectively treated as a single "analyte present" class (label 1) to capture the presence of target DNA or viral RNA in each sample.

**Table S1. Performance Comparison of Proposed Model with Contemporary Models on Target-DNA for Analyte Detection task. Friedman ranking and Wilcoxon signed-rank test $p$-values vs our model are reported for each metric. Statistically significant $p$-values ($p < 0.05$) are shown in bold.**

| Model | Precision | | | Recall | | | F1-Score | | | AUC | | |
|---|---|---|---|---|---|---|---|---|---|---|---|---|
| | Score | Rank | $p$-val | Score | Rank | $p$-val | Score | Rank | $p$-val | Score | Rank | $p$-val |
| **Our Model (Mc-GNN)** | **0.9909** | **2.5** | – | **0.9908** | **1.1** | – | **0.9908** | **1.0** | – | **0.9974** | **1.0** | – |
| Shiaelis et al. (1) | 0.9759 | 5.6 | **0.0312** | 0.9696 | 2.5 | **0.0312** | 0.9724 | 2.2 | **0.0312** | 0.9871 | 2.0 | **0.0312** |
| ShallowCNN-1 (2) | 0.9601 | 6.4 | **0.0312** | 0.9088 | 3.8 | **0.0312** | 0.9332 | 2.8 | **0.0312** | 0.9661 | 3.0 | **0.0312** |
| ShallowCNN-2~ (3–5) | 0.9880 | 3.1 | 0.5000 | 0.5152 | 7.6 | **0.0312** | 0.6752 | 7.6 | **0.0312** | 0.6920 | 7.4 | **0.0312** |
| GhostNet (6) | 0.9690 | 4.7 | 0.2188 | 0.6232 | 6.7 | **0.0312** | 0.7512 | 6.8 | **0.0312** | 0.8323 | 4.8 | **0.0312** |
| ShuffleNet (7) | 0.9614 | 4.5 | **0.0339** | 0.6340 | 5.8 | **0.0312** | 0.7571 | 5.8 | **0.0312** | 0.7942 | 5.4 | **0.0312** |
| ResNet9 (8) | 0.9852 | 3.5 | 0.4062 | 0.4353 | 8.3 | **0.0312** | 0.5901 | 8.0 | **0.0312** | 0.5919 | 8.8 | **0.0312** |
| MobileNet (9) | 0.8732 | 6.7 | 0.0625 | 0.7635 | 4.4 | **0.0339** | 0.7849 | 5.6 | **0.0312** | 0.6555 | 7.2 | **0.0312** |
| EfficientNet (10, 11) | 0.8476 | 8.0 | **0.0312** | 0.8289 | 4.8 | **0.0312** | 0.8173 | 5.2 | **0.0312** | 0.8160 | 5.4 | **0.0312** |

Figure **??**a and **??**b show the confusion matrices for the best test folds on the Target-DNA and Viral-RNA datasets, respectively. The proposed Mc-GNN achieves accurate classification for the Target-DNA dataset in control and near-perfect separation 99.72% in analyte-present samples. For Viral RNA, appreciable classification is achieved- 95.97% controls and 97.24% analyte-present cases are correctly identified. These minimal misclassifications support the sensitivity and specificity of the approach. In binary type analyte detection tasks, in general, the recall value reported corresponds to the recall of the positive class (typically class 1), which is equivalent to its class-wise accuracy. This should not be confused with macro-averaged recall unless explicitly computed. Thus, Figure **??**c presents the comparative average class-wise accuracy for analyte detection across all models. Mc-GNN attains the highest performance, with 98.19% on the Target-DNA dataset and 95.66% on the Viral-RNA dataset. The strongest baseline, Shiaelis et al. (1), achieves 94.99% (Target-DNA) and 90.17% (Viral-RNA). Shallow-CNN-1 achieves closer to 90% in Target-DNA, but performs poorly in the other dataset. while all other popular CNN/mobile models perform considerably worse, typically below 80% on DNA and 70% on viral RNA (with some as low as 48%, which was confirmed not to be the implementation issues). These results further emphasize the robustness and generalizability of Mc-GNN, which consistently achieves superior detection of the presence of analyte even in challenging, heterogeneous biological backgrounds. The low rate of false negatives and false positives confirms that Mc-GNN is highly suited for rapid, label-free analyte screening in both synthetic and clinically relevant viral material, providing support for the subsequent concentration prediction classification.

As summarized in Tables S1 and S2, the Friedman tests revealed statistically significant differences among models across all four evaluation metrics-Precision, Recall, F1-score, and AUC for both the Target-DNA and Viral-RNA datasets in the analyte detection task. For the Target-DNA dataset, the results were: Precision ($\chi^2 = 17.86$, $p = 2.23 \times 10^{-2}$), Recall ($\chi^2 = 29.80$, $p = 2.29 \times 10^{-4}$), F1-score ($\chi^2 = 32.48$, $p = 7.6 \times 10^{-5}$), and AUC ($\chi^2 = 36.27$, $p = 1.6 \times 10^{-5}$); for the Viral-RNA dataset: Precision ($\chi^2 = 28.37$, $p = 4.08 \times 10^{-4}$), Recall ($\chi^2 = 30.13$, $p = 2.00 \times 10^{-4}$), F1-score ($\chi^2 = 34.83$, $p = 2.9 \times 10^{-5}$), and AUC ($\chi^2 = 36.59$, $p = 1.4 \times 10^{-5}$). In both datasets, Mc-GNN consistently achieved the best or near-best average ranks across all metrics (rank 1.0 on Target-DNA; top ranks in F1, AUC, and Recall, and second in Precision on Viral-RNA), demonstrating robust and balanced performance. Post-hoc Wilcoxon signed-rank tests confirmed statistically significant improvements ($p < 0.05$) of Mc-GNN over most baselines. Specifically, Mc-GNN significantly outperformed Shiaelis *et al.* (1), ShuffleNet (7), MobileNet (9), EfficientNet (10, 11), and ShallowCNN-1 (2) across all metrics. While ResNet9 (8) and ShallowCNN-2 (3–5) achieved relatively high precision, they ranked lowest in Recall, F1-score, and AUC, indicating poor generalization. Similarly, MobileNet showed strong recall but weaker precision and AUC, resulting in less balanced performance. Overall, these findings highlight the consistent superiority of Mc-GNN across both datasets, with significantly better discriminative ability (AUC) and generalization balance than competing CNN and hybrid baselines.

**Comparative Performance Analysis.** As shown in Table 1 of the main manuscript, the Friedman test revealed statistically significant differences across the evaluated baselines on the Target-DNA concentration prediction task for all three metrics: precision ($\chi^2 = 31.36$, $p = 1.21 \times 10^{-4}$), recall ($\chi^2 = 32.96$, $p = 6.3 \times 10^{-5}$), and F1-score ($\chi^2 = 33.49$, $p = 5.0 \times 10^{-5}$). Our proposed Mc-GNN consistently achieved the top rank (1.0) across all metrics, delivering precision of 0.9813, recall of 0.9818, and F1-score of 0.9814. Post-hoc Wilcoxon signed-rank tests confirmed that Mc-GNN's improvements over each baseline were statistically significant ($p = 0.0312$ for all metrics).

The strongest baseline, Shiaelis *et al.* (1), attained the second-best rank, demonstrating the capability of particle-resolved fluorescence models but relying on fluorescence labels for signal enhancement. ShallowCNN-1 (2) followed in third place with moderate accuracy, reflecting the strengths and limits of lightweight 2D convolutions. In contrast, compact mobile-optimized architectures such as ResNet9 (8), GhostNet (6), ShuffleNet (7), MobileNet (9), and EfficientNet (10, 11) showed unbalanced precision–recall trade-offs, resulting in low F1-scores ($\leq 0.35$) and ranks between 5.2 and 7.8. ShallowCNN-2 (3–5) performed poorest, confirming the limitations of deeper image-level convolutional models lacking explicit morphological or relational

**Table S2. Performance Comparison of Proposed Model with Contemporary Models on target Viral-RNA for Analyte Detection task. Friedman ranking and Wilcoxon signed-rank test $p$-values vs our model are reported for each metric. Statistically significant $p$-values ($p < 0.05$) are shown in bold and second best in underline.**

| Model | Precision | | | Recall | | | F1-Score | | | AUC | | |
|---|---|---|---|---|---|---|---|---|---|---|---|---|
| | Score | Rank | $p$-val | Score | Rank | $p$-val | Score | Rank | $p$-val | Score | Rank | $p$-val |
| **Our Model (Mc-GNN)** | **0.9813** | 1.8 | – | **0.9516** | **2.0** | – | **0.9661** | **1.0** | – | **0.9916** | **1.0** | – |
| Shiaelis et al. (1) | 0.9525 | 3.4 | **0.0312** | 0.9001 | 3.4 | **0.0312** | 0.9253 | 2.0 | **0.0312** | 0.9726 | 2.0 | **0.0312** |
| ShallowCNN-1 (2) | 0.7996 | 5.4 | **0.0312** | 0.8287 | 3.6 | 0.1562 | 0.8021 | 4.0 | **0.0312** | 0.7684 | 3.2 | **0.0312** |
| ShallowCNN-2~ (3–5) | 0.8111 | 5.8 | **0.0312** | 0.2884 | 8.0 | **0.0312** | 0.3956 | 8.0 | **0.0312** | 0.4777 | 7.4 | **0.0312** |
| GhostNet (6) | 0.7222 | 7.2 | 0.0625 | 0.4191 | 6.8 | **0.0312** | 0.4378 | 6.8 | **0.0312** | 0.4684 | 7.4 | **0.0312** |
| ShuffleNet (7) | 0.7603 | 5.8 | **0.0312** | 0.6636 | 5.8 | **0.0312** | 0.7019 | 5.6 | **0.0312** | 0.6136 | 5.0 | **0.0312** |
| ResNet9 (8) | 0.9806 | **1.6** | 0.5000 | 0.1590 | 8.6 | **0.0312** | 0.2713 | 8.6 | **0.0312** | 0.4426 | 8.4 | **0.0312** |
| MobileNet (9) | 0.6773 | 8.4 | **0.0312** | 0.9262 | 2.8 | 0.3125 | 0.7819 | 4.8 | **0.0312** | 0.5279 | 6.6 | **0.0312** |
| EfficientNet (10, 11) | 0.7642 | 5.6 | **0.0312** | 0.8444 | 4.0 | 0.0938 | 0.7971 | 4.2 | **0.0312** | 0.7172 | 4.0 | **0.0312** |

**Table S3. Ablation study of the proposed Mc-GNN on both datasets for the concentration-prediction task. Average 5-fold scores, Friedman rankings, and Wilcoxon signed-rank $p$-values (vs. Mc-GNN) are reported. Statistically significant differences ($p < 0.05$) are shown in bold.**

**Target-DNA Dataset**

| Model | Precision | | | Recall | | | F1-Score | | |
|---|---|---|---|---|---|---|---|---|---|
| | **Score** | **Rank** | $p$-val | **Score** | **Rank** | $p$-val | **Score** | **Rank** | $p$-val |
| **Mc-GNN** | **0.9813** | **1.0** | – | **0.9818** | **1.0** | – | **0.9814** | **1.0** | – |
| Morphology-Guided Conv. | 0.9636 | 2.0 | **0.0312** | 0.9663 | 2.0 | **0.0312** | 0.9643 | 2.0 | **0.0312** |
| Fixed-Shape Kernels + GNN | 0.9069 | 3.4 | **0.0312** | 0.9206 | 3.0 | **0.0312** | 0.9103 | 3.2 | **0.0312** |
| Fixed-Shape Kernel Conv. | 0.8805 | 3.6 | **0.0312** | 0.8799 | 4.0 | **0.0312** | 0.8729 | 3.8 | **0.0312** |

**Viral-RNA Dataset**

| Model | Precision | | | Recall | | | F1-Score | | |
|---|---|---|---|---|---|---|---|---|---|
| | **Score** | **Rank** | $p$-val | **Score** | **Rank** | $p$-val | **Score** | **Rank** | $p$-val |
| **Mc-GNN** | **0.9476** | **1.0** | – | **0.9480** | **1.0** | – | **0.9474** | **1.0** | – |
| Morphology-Guided Conv. | 0.9302 | 2.2 | **0.0312** | 0.9266 | 2.2 | **0.0312** | 0.9279 | 2.2 | **0.0312** |
| Fixed-Shape Kernels + GNN | 0.8646 | 2.8 | **0.0312** | 0.8730 | 2.8 | **0.0312** | 0.8646 | 2.8 | **0.0312** |
| Fixed-Shape Kernel Conv. | 0.7039 | 4.0 | **0.0312** | 0.7212 | 4.0 | **0.0312** | 0.6953 | 4.0 | **0.0312** |

modeling.

Consistent results were observed for the Viral-RNA dataset as shown in the Table 2 of the main manuscript , where the Friedman test also detected significant performance differences across baselines-precision ($\chi^2 = 28.53$, $p = 3.83 \times 10^{-4}$), recall ($\chi^2 = 30.88$, $p = 1.48 \times 10^{-4}$), and F1-score ($\chi^2 = 34.19$, $p = 3.80 \times 10^{-5}$). Mc-GNN again achieved the top rank (1.0) across all metrics, with precision, recall, and F1-scores of 0.9476, 0.9480, and 0.9470, respectively, and all Wilcoxon post-hoc tests against baselines reaching statistical significance ($p = 0.0312$). Shiaelis *et al.* retained the second rank, while ShallowCNN-1 ranked third but failed to capture complex spatial–morphological clustering patterns. MobileNet and EfficientNet yielded balanced recall but low precision (F1 $< 0.20$), whereas ShuffleNet, GhostNet, and ResNet9 exhibited uneven or unstable trade-offs. Across both datasets, these consistent findings reinforce the effectiveness of Mc-GNN's morphology-guided dual-stage architecture in capturing particle-level structure–function relationships.

**Ablation Study.** To evaluate the contribution of individual architectural components in our proposed Mc-GNN model, we conducted an ablation study on the concentration prediction task using both the Target-DNA and Viral-RNA datasets. The baseline variants included: *(i)* Morphology-Guided Convolution, *(ii)* Fixed-Shape Kernel Convolution, and *(iii)* Fixed-Shape Kernels with GNN. The ablation study focused on the concentration prediction task, which forms the primary focus.

We evaluated all models across five stratified folds and reported the mean Precision, Recall, and F1-score along with Friedman rankings and Wilcoxon signed-rank test results against the full Mc-GNN model (see Table S3). Friedman tests confirmed statistically significant differences in model performance across all three metrics on both datasets: Target-DNA dataset: Precision ($\chi^2 = 13.56$, $p = 0.0036$), Recall ($\chi^2 = 15.00$, $p = 0.0018$), and F1-score ($\chi^2 = 14.04$, $p = 0.0029$); Viral-RNA dataset:

Precision, Recall, and F1-score all yielded identical values ($\chi^2 = 14.04$, $p = 0.0029$). Across both datasets and all metrics, Mc-GNN achieved the best average rank (1.0), demonstrating the synergistic benefit of combining morphology-guided kernels and graph-neural network. Post-hoc Wilcoxon signed-rank tests revealed that Mc-GNN significantly ($p < 0.05$) outperformed all ablation variants across every metric, underscoring the importance of each design choice. Notably, obtaining the best rank in four out of five folds, but a lower rank or tie in the remaining fold, can yield a moderate yet statistically significant $p$-value (e.g., $p = 0.0312$). Such a scenario illustrates that even small but consistent improvements, accompanied by occasional ties or minor rank variations, can result in statistically meaningful outcomes in Wilcoxon signed-rank tests.

**Table S4. ssDNA Sequences for Target-DNA and Thiolated Probes**

| ssDNA Strand ID | Sequence (5' to 3') |
|---|---|
| E-DNA (from SARS-CoV-2 *E* Gene; NCBI `NC_045512.2`) | agagacaggtacgttaatagttaatagcgtacttctttttcttgctttcgtggtattctt gctagttacactagccatccttactgcgcttcgattgtgtgcgtactgctgcaatattgtt aacgtgagtcttgtaaaaccttctttttacgtttactctcgtgttaaaaatctgaattctt ctagag |
| N-DNA (from SARS-CoV-2 *N* Gene; NCBI `NC_045512.2`) | actcaacatggcaaggaagaccttaaattccctcgaggacaaggcgttccaattaacacc aatagcagtccagatgaccaaattggctactaccgaagagctaccagacgaattcgtggtg gtgacggtaaaatgaaagatctcagtccaagatggtatttctactacctaggaactgggcc agaagctggacttcccta |
| PrE1 | SH-TTTTTTTTTT-aactattaacgtacctgtctct |
| PrE2 | SH-TTTTTTTTTT-ctctagaagaattcagattt |
| PrN1 | SH-TTTTTTTTTT-aatttaaggtcttccttgcc |
| PrN2 | SH-TTTTTTTTTT-tagggaagtccagcttctgg |

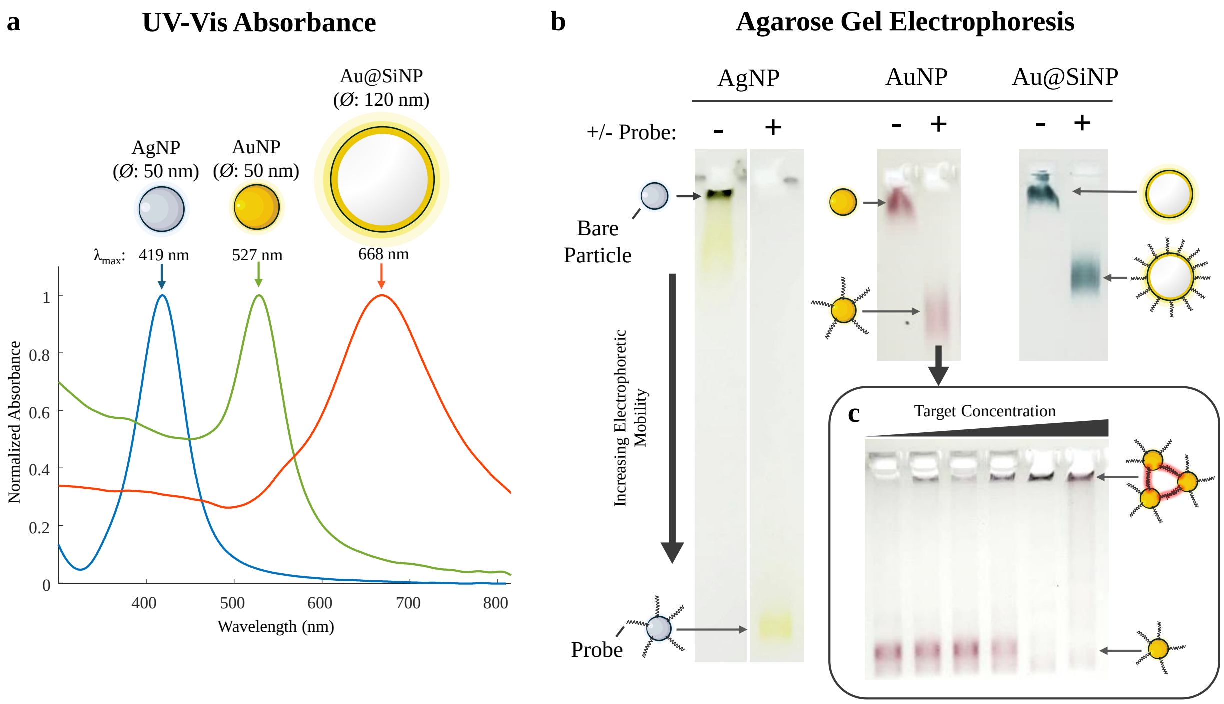


**Fig. S2.** Nanoparticles with unique spectral properties are conjugated with DNA for use in the assay. a) Normalized UV visible absorbance spectra for the different particles in use, demonstrating their different scattering properties. Spectra for 50 nm diameter spherical silver nanoparticles (AgNPs), 50 nm diameter spherical gold nanoparticles (AuNPs), and 120 nm diameter gold-shelled silica-core particles (Au@SiNPs) are shown. Arrows indicate the wavelength of maximum of scattered light for each particle type. b) Agarose gel electrophoresis is used to quickly characterize the conjugation and aggregation of particles. AgNPs were run a 0.5% agarose gel. AuNPs and Au@SiNPs were run on 1% agarose gels. Successful conjugation of probe DNA to a particle adds significant negative charge to a particle and increases its electrophoretic mobility (runs farther on the gel). If particles are aggregated, they will generally be confined to the well and not be able to move in the gel. Here, conjugated and unaggregated particles (single bands) are shown for each particle type. c) In the inset of (b), an aggregation example with gel readout is shown for Pr-E1 and Pr-E2 AuNPs (Table S3). Concentration of the target (E-DNA, Table S3) increases in each lane from left to right (0 pM, 1 pM, 10 pM, 100 pM, 1 nM, and 10 nM respectively). Even at 1 pM, aggregates are seen in the well.
